# Modeling cannabinoids from a large-scale sample of *Cannabis sativa* chemotypes

**DOI:** 10.1101/2020.02.28.970434

**Authors:** Daniela Vergara, Reggie Gaudino, Thomas Blank, Brian Keegan

## Abstract

The accelerating legalization of *Cannabis* has opened the industry to using contemporary analytical techniques. The gene regulation and pharmacokinetics of dozens of cannabinoids remain poorly understood. Because retailers in many medical and recreational jurisdictions are required to report chemical concentrations of cannabinoids, commercial laboratories have growing chemotype datasets of diverse *Cannabis* cultivars. Using a data set of 17,600 cultivars tested by Steep Hill Inc., we apply machine learning techniques to interpolate missing chemotype observations and cluster cultivars together based on similarity. Our results show that cultivars cluster based on their chemotype, and that some imputation methods work better than others at grouping these cultivars based on chemotypic identity. However, due to the missing data for some of the cannabinoids their behavior could not be accurately predicted. These findings have implications for characterizing complex interactions in cannabinoid biosynthesis and improving phenotypical classification of *Cannabis* cultivars.

## Introduction

The legalization of medicinal and recreational *Cannabis* consumption represents one of the most dramatic shifts in social policy since same-sex marriage (1). Multiple European and South American countries have legalized its medicinal and recreational use. In North America, Mexico has legalized its medicinal use, Canada permits both medicinal and recreational consumption, and in the US, it varies on a state-by-state basis. As of August 2018, nine states and the District of Columbia legalized the medical and recreational use of *Cannabis*, while 29 other states legalized its medical consumption. *Cannabis* has opened the doors to a new—and highly-profitable—worldwide industry, with medical and agricultural research potential. In the first year after legalization in Colorado, sales of *Cannabis* totaled over $700 million USD and by 2017 cumulative sales surpassed $4 billion. In Canada, the estimated value of the marijuana market by 2020 will be around CAD$6.8 billion CAD (approximately USD$4.5 billion) (https://daily.jstor.org how-big-will-canadas-legal-cannabis-market-be).

*Cannabis*, is an angiosperm (flowering) plant, which produces the cannabinoid compounds driving this growing market. Cannabinoids are a specific type of terpenoid that include approximately 25,000 different chemicals (2), some of which can interact with the human endocannabinoid system within the brain and nervous system. *Cannabis* produces as many as 120 different cannabinoids, some of which have demonstrated psychoactive effects and medicinal potential (3). These compounds are most abundant in the trichomes of female flowers (4). The plant produces these compounds in the acidic form, and are reduced into the neutral form, typically by heating. Two of the most known cannabinoids, Δ-9-tetrahydrocannabinolic acid (THCA) and cannabidiolic acid (CBDA), are converted respectively to their neutral forms Δ-9 tetrahydrocannabinol (THC) and cannabidiol (CBD), when heated. Both THCA and CBDA are mostly similar in their biochemistry, but the neutral form, THC, is well-known for its psychoactive effects (3). Growers have used a combination of selective breeding and other agricultural practices to increase the quantity of THCA produced by *Cannabis* plants over time (5, 6).

The biochemical pathway responsible for the production of these compounds is complex and involves multiple genes (7). The enzymes responsible for the production of CBDA and THCA—CBDA and THCA synthases, respectively—act on the same precursor molecule: Cannabigerolic acid (CBGA) (8). Therefore, if the plant has THCA synthase it produces THCA, if it has CBDA synthase it will produce CBDA, and if it has both synthases it will produce both compounds. These two synthases are found at the final stage of the biochemical pathway, along with a third poorly studied synthase, Cannabichromenic acid synthase, that produces a third compound, Cannabichomenic acid (CBCA). The genes responsible for these enzymes are very similar in their genetic sequence (7) and are very close in proximity (9–11), suggesting that they might have originated from the same ancestor gene (12). Given that these synthases act on the same precursor molecule and that their DNA sequence is so similar, they could be considered “promiscuous enzymes” (13–15), and due to their vast similarities, CBDA synthase might produce some THCA and vice versa. The complexities in these synthesis pathways complicate efforts to predict or design fundamental phenotypic characteristics like THCA production in a given strain.

Other minor cannabinoids include Cannabinol (CBN), Δ-9-tetrahydrocannabivarin carboxylic acid (THCVA), and Cannabidivarinic Acid (CBDVA). CBN is a product that accumulates with the breakdown of THC (16–18), usually when *Cannabis* flowers are stored under poor conditions including high heat and long time periods (16–19). Studies imply that similar to THCA and CBDA, through a decarboxylation process with heat, THCVA and CBDVA form Δ-9-tetrahydrocannabivarin (THCV), and cannabidivarin (CBDV), respectively (20). Both THCV and CBDV are produced in very low quantities by *Cannabis* plants, and therefore have been poorly studied. Some studies suggest that both THCV and CBDV are produced by a variant of the well-known cannabinoid pathway (20, 21) that yields to the production of homologous compounds such as cannabigerovarinic acid (CBGVA) which is the precursor molecule used by THCVA and CBDVA synthases (20, 21). Even though THCV appears to produce similar psychoactive effects to those of THC, also interacts with the CB1 and CB2 endocannabinoid receptors, and may have medical use (22), the exact biochemical pathway used by the plant to produce this compound remains unexplored. In the same manner, CBDV is also poorly investigated despite it’s possible medicinal properties as an anticonvulsant (23).

The number of species that compose this genus *Cannabis* is debated among scientists: some claim that it is composed of at least two species (*C. sativa* and *C. indica*) (24), while others claim that it is composed of one species (*C. sativa*) (25). The phenotypic characteristics of these different groupings—mainly called “indica” and “sativa” within the *Cannabis* industry—including the cannabinoid production, are an important factor for the plant’s classification. “Sativa” type plants are reputed to produce high THCA and low CBDA, and are described as having uplifting and stimulating psychoactive effects after consumption. “Indica” type plants on the other hand, are reputed to have relaxing sedative effects perhaps due to its similar ratio of THCA and CBDA production (26, 27). The crosses between “sativa” and “indica” plants are referred to as “hybrids” and these have variable phenotypes usually intermediate to the parents (26). These distinctions are of critical importance particularly for medical patients, given that “sativa” plants are utilized to treat depression, low energy, headaches, nausea, and loss of appetite, while “indica” plants are typically used to treat depression, anxiety, insomnia, pain, inflammation, muscle spasms, epilepsy, and glaucoma (27). The taxonomic origins of the sativa and indica descriptions do not address the best use in medical treatment.

The taxonomic based naming convention from the *Cannabis* industry is flawed given that varieties and groupings are often mislabeled and their nomenclature does not agree with biological relatedness or therapeutic uses between plant types (26, 28, 29). For example, plants that are purportedly “sativa” and even plants that are supposed to be from the same variety (*i.e*. Girl Scout Cookies) might be very distantly related and thus have significantly different chemical profiles, while two plants that are supposed to be “indica” and “sativa” might be very closely related (26, 28, 29). This misleading identification system is particularly problematic for medical patients that are looking for reliability and consistency in their treatments. This problem is exacerbated by the fact that scientists and medical professionals in the US are not allowed to research the *Cannabis* produced by the industry because of federal regulations. Instead, *Cannabis* researchers can only research the plant material produced under license from the federal government, which poorly represents the quality and diversity currently produced by medicinal and recreational growers and breeders (19, 30).

Given this flaw in the industry with *Cannabis* variety names, a possible way to categorize and understand the relationship between different varieties is through cluster analyses. Cluster analysis groups data points whose identities are unknown, allowing for varieties with similar cannabinoid profiles to group together. One approach to clustering commonly used in machine learning involves dimensionality reduction, where a highdimensional representation of data is transformed into a lower-dimensional representation while preserving as much variance as possible (31). Although Principal Component Analysis is the most used dimensionality reduction method in biology, there are other multiple tools for this purpose such as t-distributed stochastic neighbor embedding (t-SNE) (32) and uniform manifold approximation and projection (UMAP) (33). Like PCA, these dimensionality reduction methods can take a high-dimensional vector (*i.e.*, multiple observations of chemical concentrations) and reduce it to a low-dimensional vector (*i.e.*, two dimensions for visualization and clustering), but can capture clusters or preserve more of the variance with which linear methods like PCA might struggle in a low dimensional visual representation.

## Methods

### Data collection

Cannabinoid concentration profiles (chemotypes) were generated by Steep Hill, Inc following their published protocol (29). Briefly, data collection was performed using high performance liquid chromatography (HPLC) with Agilent (1260 Infinity, Santa Clara, CA) and Shimadzu (Prominence HPLC, Columbia, MD) equipment with 400-6000 mg of sample. We only used data from *Cannabis* flowers and exclude those from concentrates and edibles.

We combined the acidic forms of the cannabinoids and their neutral forms using the same conversion rate of 0.88 which was also used by Vergara et al. (2017). The combination of the acidic and neutral forms provides a more honest analysis given that these flowers could have been collected at different points in time and therefore their ratio of acidic to neutral forms of the cannabinoids could differ. We analyzed seven cannabinoids, four which are part of the same biochemical pathway (7): CBG, CBD, THC, CBC and the breakdown product of THC, Cannabinol (CBN). We also included for some of our analyses two other cannabinoids whose biochemical pathways is still unknown: Tetrahydrocannabivarin (THCV) and Cannabidivarin (CBDV).

### Data distribution

The raw data provided by Steep Hill, Inc. consisted of 13 different files and is found in the dryad repository (link). These data were taken from a combination of regulatory compliance testing, internal customer research and development, and data collected as part of studies directed to chemical or genetic characterization. Therefore, these data represent a mixture of types that were combined and subsequently filtered based on depth of information. We used custom Python scripts using the pandas library to clean these raw files, which resulted in a single file with 17,600 entries, with samples from four different US states: 2 from Alaska; 16,418 from California; 936 from Colorado; and 244 from Washington. California was the only state that included samples from eight years (2011–2018), and the only year that included three states (California, Colorado, and Washington) was 2014.

### Missing data and imputation

Few of these cultivars (*N*=153) have information about all of the major cannabinoids summarized in table 1 with the features of the *N*=17,600 unique cultivars in the data set. Given the overwhelming prevalence of cases missing at least one observation of a cannabinoid (table 1, Figure S1), dropping these cases from the analysis would greatly reduce statistical power as well as introduce significant sampling biases. We follow statistical best practices about reporting missing data by reporting on the completeness of the data, detail our approaches for handling missing data, and explore the features of the missing data.

**Table 1.** Cannabinoid distribution for 17,612 unique cultivars. For each cannabinoid (column 1), we report the number of cases (column 2), the percentage missing (column 3), the average concentration (column 4), standard deviation of concentrations (column 5), and maximum observed value (column 8).

| **Cannabinoid** | **Count** | **Missing** | **Mean** | **S.D.** | **Max.** |
| --- | --- | --- | --- | --- | --- |
| Cannabigerol (CBG) | 12,270 | 30.28% | 0.50% | 0.47% | 11.58% |
| $\Delta^9$-tetrahydrocannabinol (THC) | 17,346 | 1.44% | 14.03% | 6.45% | 33.95% |
| Cannabidiol (CBD) | 10,948 | 37.80% | 1.68% | 3.91% | 25.80% |
| Cannabichromene (CBC) | 6,487 | 63.14% | 0.120% | 0.128% | 1.79% |
| Cannabinol (CBN) | 6,675 | 62.07% | 0.067% | 0.18% | 2.95% |
| Tetrahydrocannabivarin (THCV) | 3,311 | 81.19% | 0.198% | 0.412% | 6.62% |

| Cannabidivarin (CBDV) | 597 | 96.60% | 0.069% | 0.11% | 0.83% |
| --- | --- | --- | --- | --- | --- |

These data could be missing for a variety of reasons. Mechanisms of missingness include “missing completely at random” (MCAR), “missing at random” (MAR), and “missing not at random” (MNAR) (34). Some clients may elect not to test for more cannabinoids than regulations require (typically only THCA and THC) while others test for complete profiles for research, product development, or marketing purposes. The complex regulation of cannabinoid synthesis (7) and the significant genetic diversity in *Cannabis* (7, 28, 29) complicate efforts to model the relationships among cannabinoids. Methods like flux balance analysis could be used to augment the data, however the required constraints, objective functions, and metabolic pathways have not yet been well-characterized in *Cannabis.*

We instead employ imputation-based methods to estimate the missing data values. Imputation specifically refers to a class of methods for estimating missing values in a data set. The central intuition behind imputation is that information like **1)** other cannabinoids in the observation, **2)** the distribution of observed cannabinoids across samples, and **3)** similarities to other observations with complete data can guide estimation of the missing values. We employ four distinct imputation methods—iterative, multiple, k-neighbors, and soft—to estimate the missing values in our dataset using the “fancyimpute” package in Python (https://github.com/iskandr/fancyimpute). The use of these imputation methods are theoretically defensible because many of these cannabinoids are part of the same biochemical pathways and thus their prevalence and concentration are strongly correlated rather than independent (8). For example, CBGA is the precursor molecule to which THCA, CBDA, and CBCA synthases act on (7, 8), so we expect that the abundance of THCA, CBDA, and CBCA could be used to estimate the abundance of CBGA. We evaluate the performance of these imputation methods using several approaches: examining the covariance among cannabinoids, the deviance of the imputed distributions from the non-missing data, the performance of different classes of regression models, and finally visualizing the overlap of the imputed values with existing clusters of data using dimensionality reduction techniques.

### Dimensionality reduction and clustering

Dimensionality reduction are methods for transforming high-dimensional data like our seven cannabinoid observations per cultivar into a meaningful lower dimensional representation. Dimensionality reduction can be a powerful tool for classification, visualization, and compression of high-dimensional data, but have traditionally been limited to linear techniques such as principal components analysis (PCA) or factor analysis (31). However, many kinds of real-world data, like the cannabinoid chemotypes in the present study, have complex and nonlinear relationships that require alternative dimensionality reduction techniques. We apply two recently-developed and non-linear dimensionality reduction techniques, t-distributed stochastic neighbor embedding (t-SNE) (32) and uniform manifold approximation and projection (UMAP) (33) to project the seven-dimensional data down into two dimensions for the purposes of visualization and clustering. Unlike PCA, the resulting X and Y axes in t-SNE and UMAP do not have substantive interpretations of some latent dimension but their spatial proximity still captures similarity: closer points are more similar than more distant points. These non-linear projections are transductive (making sense from the observations themselves) rather than inductive (premises provide general rules), so withholding a subset of the data (*e.g.*, the imputed values) will return a different projection than using the full set. These dimensionality reduction techniques were applied to the data using the scikit-learn and umap libraries in Python (33).

### Statistical modeling

To evaluate the performance of the imputed data, we treated each cannabinoid as a dependent variable in a linear equation with the other cannabinoids as explanatory variables. This allow us to use several different regression methods to estimate the fit of the model to the data. Regression models are drawn from the traditional ordinary least square linear regression as well as more advanced k-neighbors and support vector regression, which can model non-linear relationships. Models are then fitted using data from each of the four different imputation methods. The performance of each model is evaluated using the coefficient of determination (R^2^), where values closer to 0 indicate poor fit and values closer to 1 indicate perfect fit.

## Results

### Missing data and imputation

Our data shows substantial evidence of missing data which is not at random, likely due to regulatory requirements, market demand, and client preferences. Regulatory requirements for chemotype testing vary over time and across jurisdictions but typically require testing for the psychoactive cannabinoid THC. These testing requirements explains the low level of missing information (1.4 %) for THC in our data (Table S1, Figure S1). The growing popular interest in the therapeutic properties of (CBD) likewise explains its relatively low levels (37.78 %) of missing data since it influences market demand. Conversely, cannabichromene (CBC), cannabinol (CBN), tetrahydrocannabivarin (THCV), and cannabidivarin (CBDV), show much higher levels of missing data since clients elect not to test for these poorly-understood and less-popular cannabinoids.

To understand how the imputed values compared to the observed values, we plotted the kernel density estimates (Figure 1) for six of the seven cannabinoids. We excluded THC because of few missing values. The graphs, particularly for the soft imputation in purple, becomes noisier when more data is missing. Therefore, for CBG and CBD (Figures 1A and 1B) the curves are much smoother given that these cannabinoids have only 30.27% and 37.78% of the data missing respectively (Table S1). However, for CBN and THCV (Figures 1D and 1E) the curves are jagged as they are missing 62% and 81.19% of the data respectively. The most extreme case is CBDV that lacks 96.6% of the data (Figure 1F).

**Figure 1.**
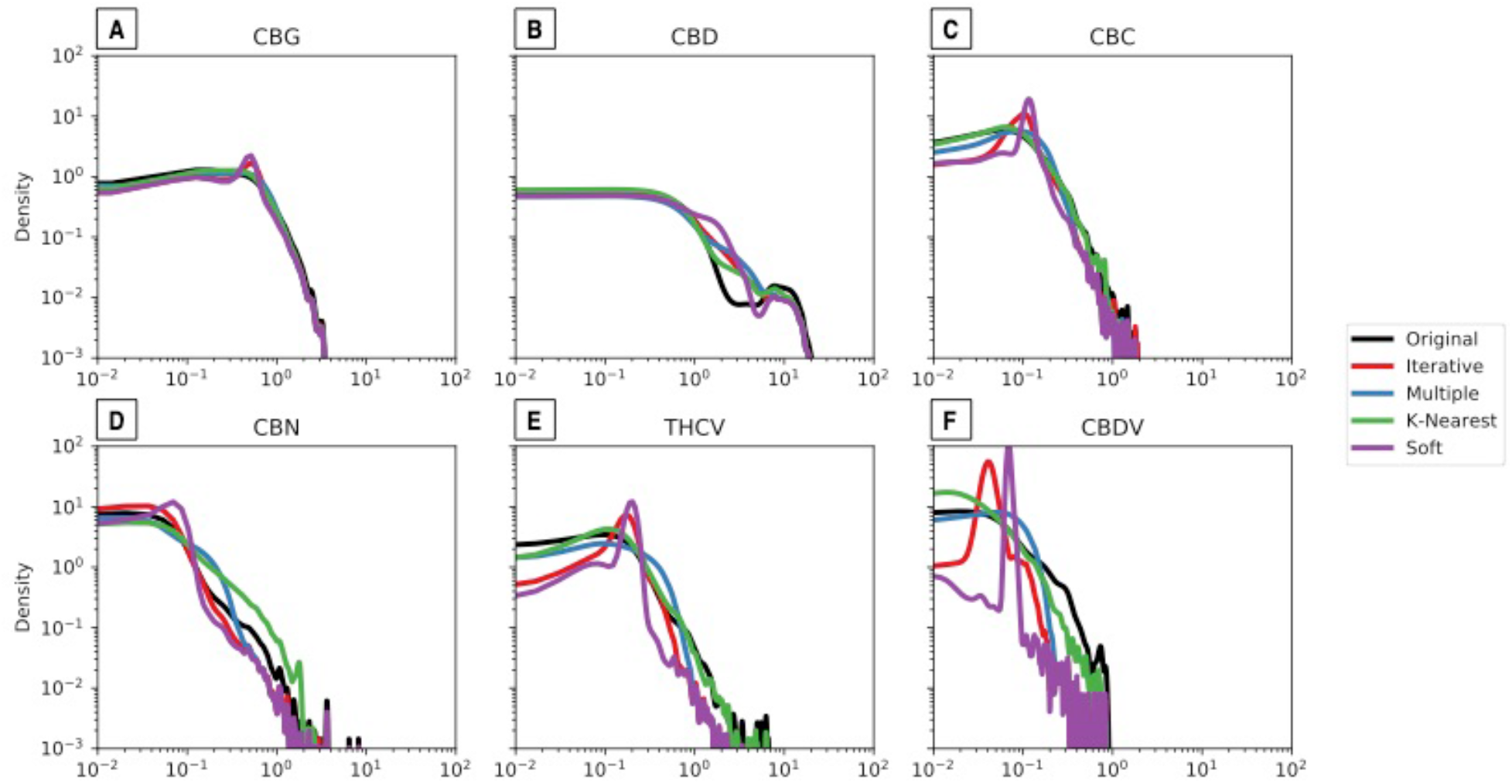
Density plots of original data and imputed values for each of the cannabinoids. The distribution of the observed values is in black, the iterative imputation in blue, the multiple imputation in red, the k-nearest neighbors in green, and the soft imputation in purple

The results of t-tests of each cannabinoid’s observed distribution against each of its imputed distributions as a measure of goodness-of-fit is summarized in Table 2. Well-fit imputed values should have small test statistics and non-significant p-values while poorly-fit imputed values will have larger test statistics and significant p-values. The imputation methods generally struggled with the CBD, CBDV, and CBN values. At first glance this under-performance could be attributed to a lack of baseline information: CBDV had missing observations in 96.6 % of the cases data and CBN was missing 60.3 %. However, THCV (81.19 %) and CBC (63.13 %) were also missing large fractions of their observations but the imputation methods returned well-fit distributions. The iterative and k-nearest neighbor imputation method had significantly different distributions for four of the seven cannabinoids while the soft imputation method had well-fit distributions across all the cannabinoids.

**Table 2.**
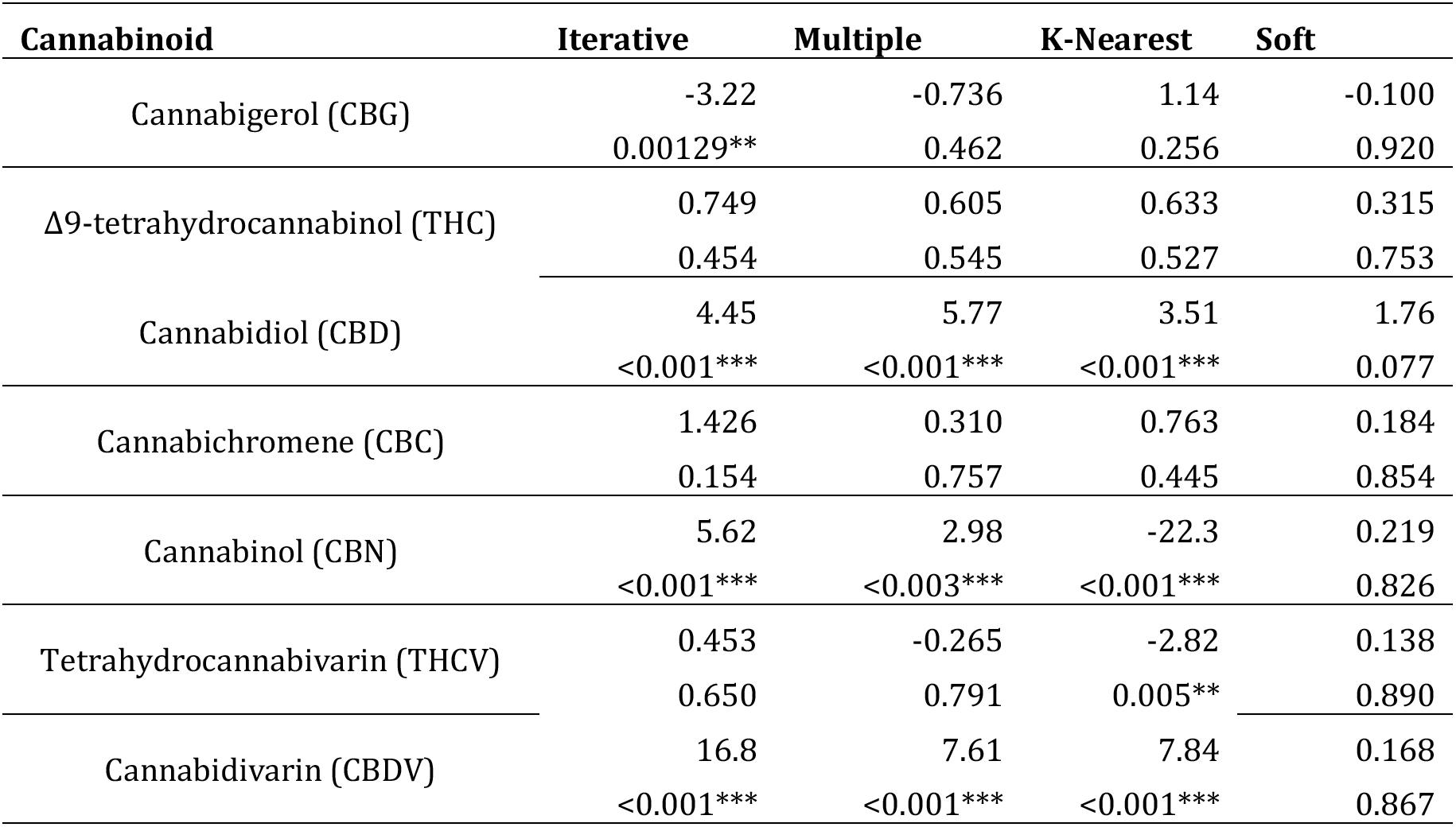
Statistics for observed vs. imputed cannabinoid concentrations. t-statistics (top) and p-values (bottom) *p < 0.05, ** p < 0.01, *** p < 0.001.

| Cannabinoid | Iterative | Multiple | K-Nearest | Soft |
| --- | --- | --- | --- | --- |
| Cannabigerol (CBG) | -3.22 | -0.736 | 1.14 | -0.100 |
|  | 0.00129** | 0.462 | 0.256 | 0.920 |
| $\Delta^9$-tetrahydrocannabinol (THC) | 0.749 | 0.605 | 0.633 | 0.315 |
|  | 0.454 | 0.545 | 0.527 | 0.753 |
| Cannabidiol (CBD) | 4.45 | 5.77 | 3.51 | 1.76 |
|  | <0.001*** | <0.001*** | <0.001*** | 0.077 |
| Cannabichromene (CBC) | 1.426 | 0.310 | 0.763 | 0.184 |
|  | 0.154 | 0.757 | 0.445 | 0.854 |
| Cannabinol (CBN) | 5.62 | 2.98 | -22.3 | 0.219 |
|  | <0.001*** | <0.003*** | <0.001*** | 0.826 |
| Tetrahydrocannabivarin (THCV) | 0.453 | -0.265 | -2.82 | 0.138 |
|  | 0.650 | 0.791 | 0.005** | 0.890 |
| Cannabidivarin (CBDV) | 16.8 | 7.61 | 7.84 | 0.168 |
|  | <0.001*** | <0.001*** | <0.001*** | 0.867 |

To understand the behavior of each of the seven cannabinoids against each other, we plotted the distribution and the bivariate relationships among the observed data (Figure 2). The performance of each cannabinoid is portrayed by the kernel-density estimate of the (log-normalized) distribution of concentrations on the diagonal (Figure 2), while the upper triangle is a scatter plot of the mutual relationships between each of the seven cannabinoids against one another. The bottom triangle are density plots of the bivariate data distributions, which show distinct clusters on the common relationship between each two cannabinoids.

**Figure 2.**
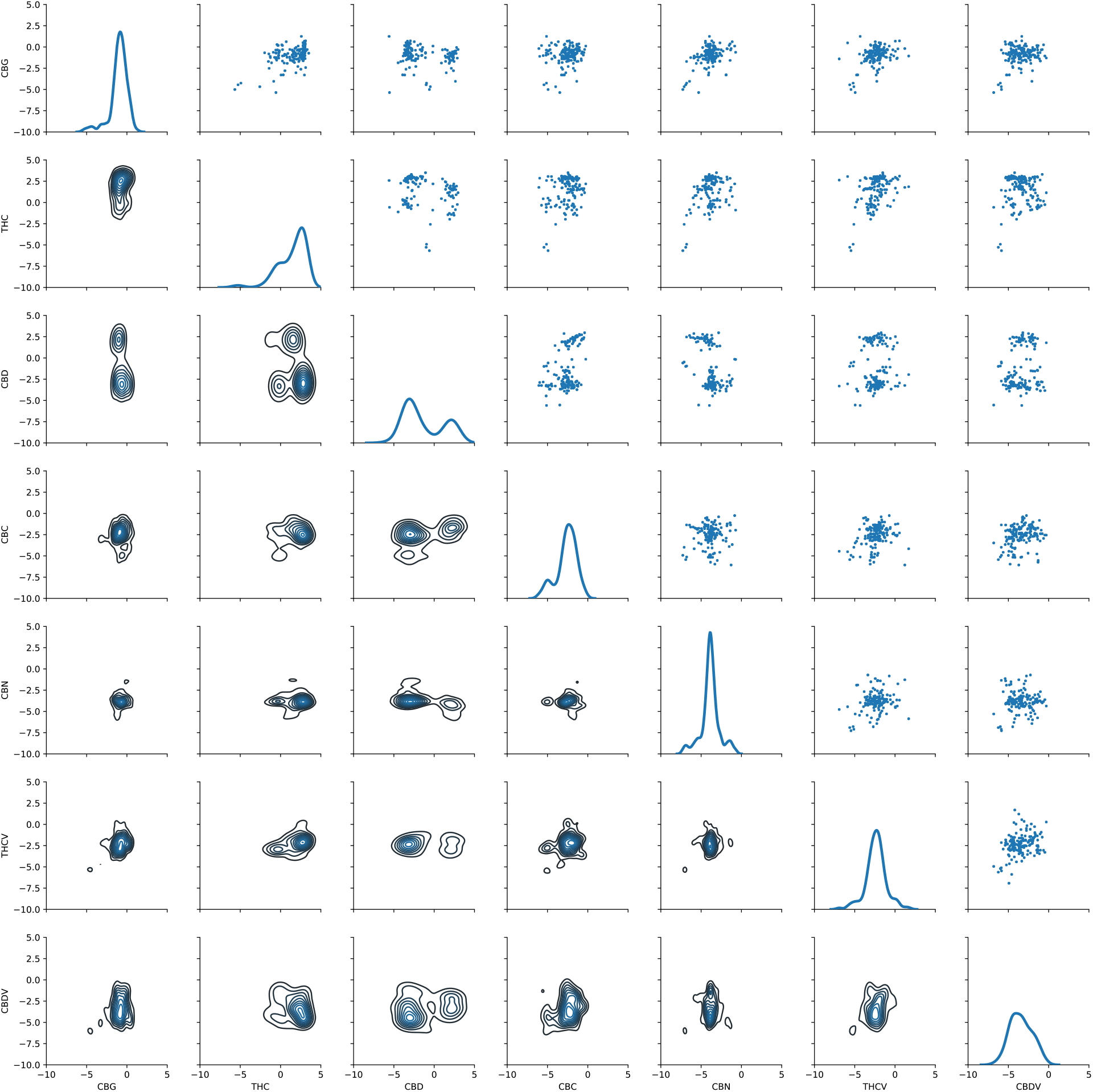
Pair plot of the observed distributions of (log-scaled) cannabinoids. Diagonal are kernel density estimates. Lower triangle are density plots and the upper triangle are scatter plots of the bivariate relationships.

The diagonal on figure 2 shows that CBG is always around 0 given that it’s found in very low quantities. On the contrary, the THC curve has two humps, a lower one around zero and a higher one in the positive side of the axis, showing that there are multiple individuals that have higher percentages of this cannabinoid. Similarly, the CBD curve also has two humps but much more conspicuous than the humps in the THC curve. The smoothness in the THC curve suggests that the data is continuous, different to CBD which has two clearly distinct groups. The bigger hump in CBD’s curve suggests that there are more individuals that have little of the cannabinoid compared to the smaller hump with individuals that produce more CBD. Even though CBC also shows two humps, both suggest that most individuals have very little CBC. In essence for CBC, CBN, THCV, and CBDV the humps have a maximum around zero establishing the lack of data for all of these cannabinoids.

The upper triangle on figure 2 shows that CBG is found in very low concentrations compared to THC, CBD, and CBC. THC is always found in the positive side of the axis indicating both that the abundance of the data for this cannabinoid and that individuals produce it in larger amounts. Finding low levels of CBG is expected given that CBGA is the precursor molecule to which THCA, CBDA, and CBCA synthases act on to convert to THCA, CBDA, and CBCA respectively. Therefore, if these synthases are active, there should be little CBGA. This is exactly what our results show. Our results also show that the three cannabinoids can coexist, THC and CBD can be present together, as well as CBC and THC and CBC and CBD. However, CBC is always found in very low levels when compared to THC and CBD.

The density plots in the lower triangle (Figure 2) confirms that CBG is found in lower quantities compared to THC and CBD, and the lack of data for the rest of the cannabinoids. Yet the relationship between CBG and CBD is interesting, with two distinct clusters. Both clusters are centered around zero in the CBG (X) axis, showing the lack of this cannabinoid, but on the CBD (Y) axis one of the clusters shows its presence. The second cluster which is the bigger one (darker blue) suggests the lack of both CBD and CBG. The relationship between THC and CBD is also interesting with three clusters suggesting that most individuals have high THC and low CBD, but the other two clusters suggest the presence of (a) high-CBD individuals and (b) individuals that produce little of either cannabinoid (e.g., low THC and low CBD possibly hemp-type).

We plotted the two most well-known and biosynthetically related cannabinoids (THC and CBD) to better visualize their relationship (Figure 3). Four distinctive clusters are apparent in this visualization. We identified four substantive clusters of cultivars based on their distinctive combinations of CBD and THC concentrations (Figure 3). Cluster 0 (orange) contains high THC and low CBD cultivars, likely intended for recreational consumption. Cluster 1 (blue) contains low THC and high CBD cultivars, likely intended for medical consumption. Cluster 2 (pink) contains high THC and high CBD cultivars, likely intended for recreational consumption. Cluster 3 (green) contains low THC and low CBD cultivars, likely hemp intended for industrial use. The DBSCAN algorithm was not able to assign the cultivars in Cluster −1 to another cluster, so these remain outliers, but could theoretically be assigned to parent clusters through iteration, alternative clustering parameterization, or other clustering methods.

**Figure 3.**
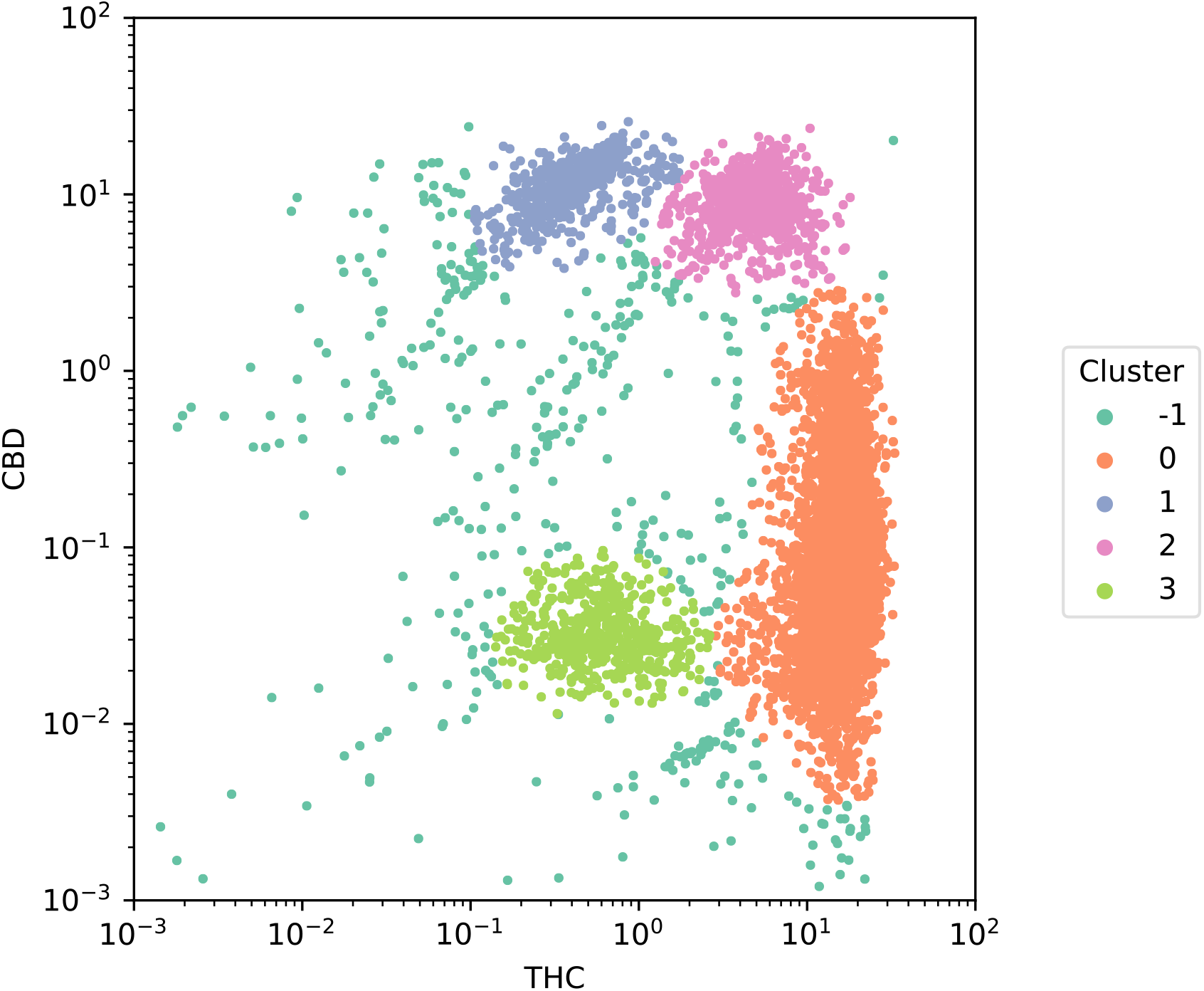
Scatter of THC against CBD concentrations colored by cluster. Four distinctive clusters are apparent in this visualization distinguished by different colors: **Cluster 0** in orange contains 8,270 cultivars and is characterized by high concentrations of THC and low concentrations of CBD, most likely intended for recreational use. **Clusters 1** in green contains 658 cultivars and is characterized by low concentrations of THC and high concentrations of CBD, most likely intended for medical consumption. **Cluster 2** in red contains 869 cultivars and is characterized by high concentrations of both THC and CBD, most likely intended for both recreational and medical use. **Cluster 3** in purple contains 536 cultivars and is characterized by low concentrations of both THC and CBD, most likely hemp-type of cultivars. Finally, **Cluster-1** in blue contains 433 cultivars that could not be assigned to other clusters using the DBSCAN algorithm, but were labeled using semi-supervised approaches like label propagation.

All of the imputation methods preserved substantive relationships like the negative THC vs. CBD correlation, and positive correlations for CBC vs. CBD, and CBN vs. CBC (Figure 4). The iterative imputation method introduced stronger correlations than existed in the original data between THC vs. CBDV, CBD vs. CBC, CBD vs. CBDV, and CBDV vs. CBC, (the red squares in the center of figure 4b). The other imputation methods preserved similar pairwise correlation structures as were found in the original data, in other words figurers S3C, D, and E are similar to figure 4A. The one that differs the most is the iterative imputation method (Figure 4b) exaggerating the relationships and displaying stronger correlations (red squares in the middle) than the original association. The K nearest neighbors (Figure 4D) is the one imputation method that weakens the relationships and the squares that are in orange in the original distribution (Fig 4A) are found in green. Notice that the relationship between THC and CBD (purple squares) in the original relationship (Fig 4A) is maintained in the other imputation methods (Figures 4B-E) oscillating between −0.5 and −0.6, suggesting a higher accuracy in predicting their behavior with any of the imputation methods. Additionally, this shows that the relationship between both cannabinoids can be easily predictable compared to any of the other cannabinoids.

**Figure 4.**
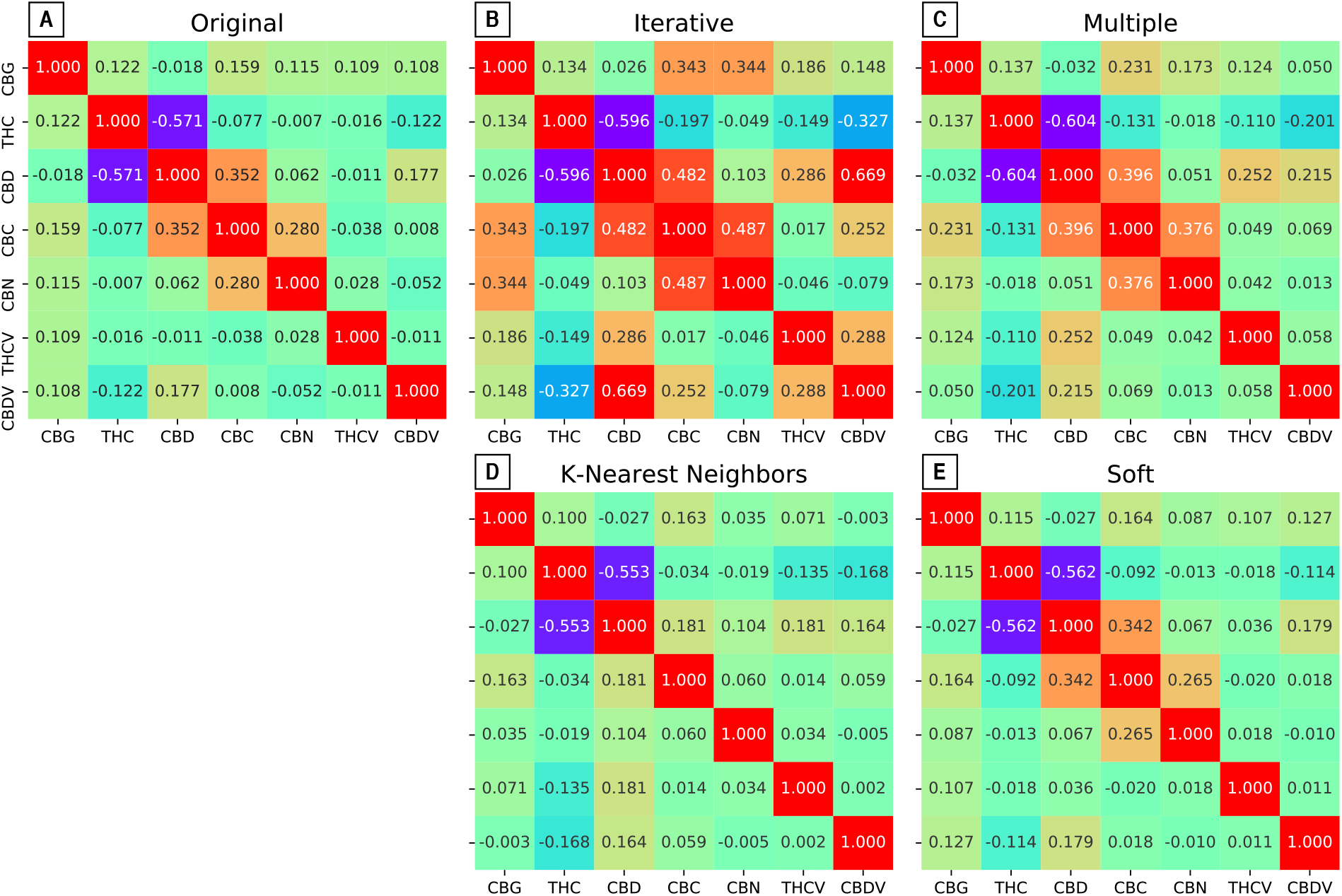
Heatmap of the symmetric pairwise Pearson correlations. Correlations were performed across the original data and the data from the four imputation methods

In order to understand how well the imputed values for the missing THC and CBD observations correspond to the clusters in figure 3, we plotted the original distribution of data (in black) and the imputed values from each of the four imputation methods (Figure 5). The iterative (Figure 5A), multiple (Figure 5B), and soft (Figure 5D) imputation methods capture a quasi-linear relationship between CBD and THC (reflecting their competition for similar precursors). The K-nearest neighbors’ method (Figure 5C) captures most of the patterns seen with the real data (Figure 3). All imputation methods capture clusters zero (high THC and low CBD), one (low THC and high CBD), and two (high THC and high CBD) from figure 3 which comprise most of the data. However, all methods overlook the underlying clustered structure and miss the low-THC, low-CBD cluster (likely hemp, cluster 3 in light green Figure 3). The failure of all methods to predict this cluster may be due to the few hemp varieties found in the data. Finally, even though K-nearest neighbor is the method that weakens the relationship between cannabinoids (Figure 4D) it is the one method that better overlays the original data for THC and CBD (Figure 5C).

**Figure 5.**
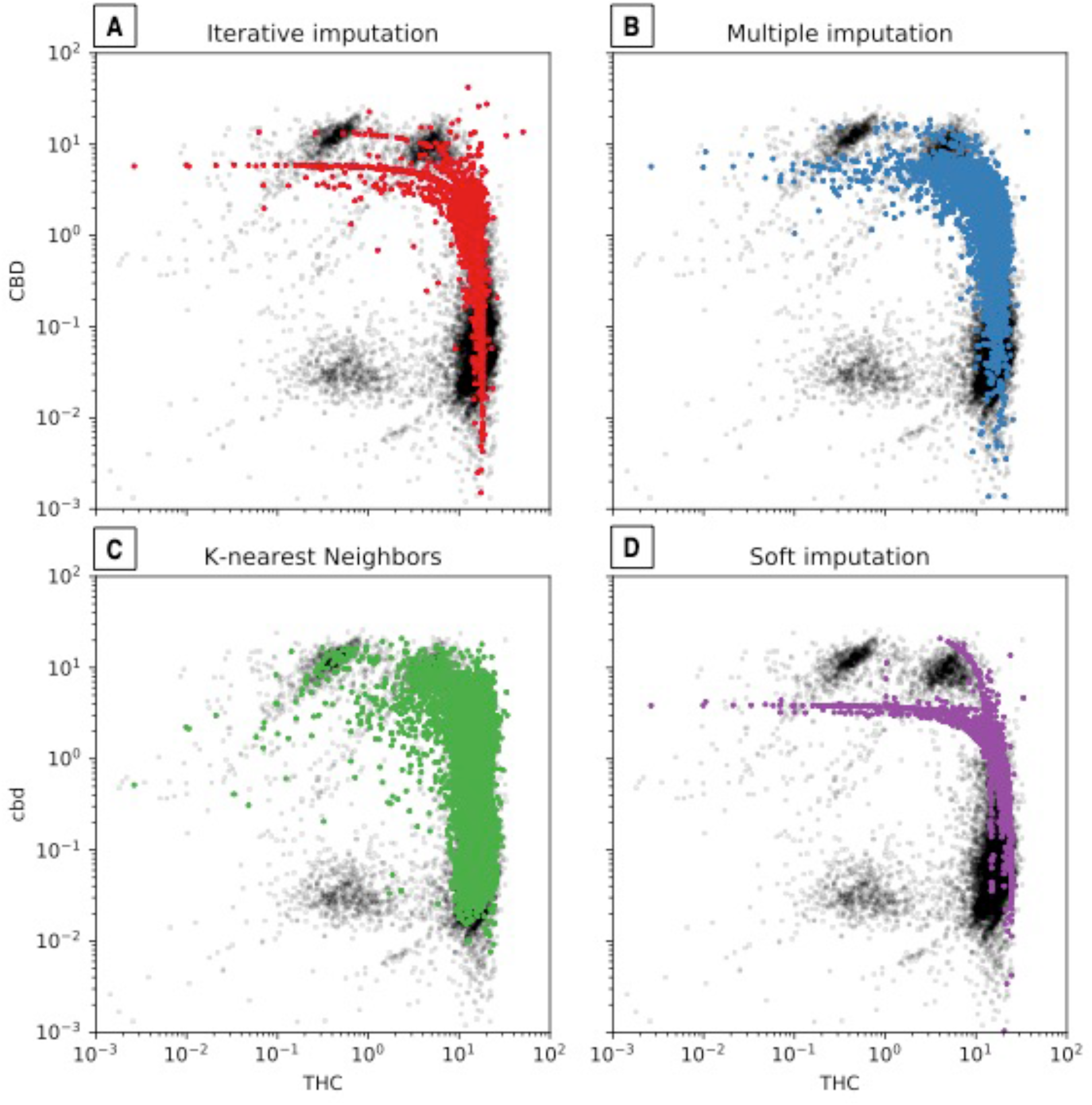
Original vs. imputed values using four methods. The 191 imputed values from each of the four methods, compared to the raw distribution on Figure 3.

### Statistical modeling

The values reported in Table 3 are the average R^2^ from the five-fold cross validation for models predicting each cannabinoid and using data from each imputation method and different regression models. Given that model performance is evaluated using five-fold cross-validation to train a regression model on a subset of 80% of the data and test its performance on a held-out set of 20% of data. As a result, Table 3 reports 12 different scores for each cannabinoid: three kinds of regression models on four kinds of imputation methods. The goal of this analysis is to identify if a particular combination of regression modeling and data imputation reliably models multiple kinds of cannabinoid abundance as a function of other cannabinoids. No regression model or imputation method is consistently better at predicting the cannabinoid concentration than others, but in some cases, particular cannabinoids can be very accurately predicted as is the case for CBD in the support vector model under the iterative imputation method (R^2^=0.815). Additionally, some cannabinoids are more accurately predicted than others, which is the case of CBD or THC compared to CBC or CBDV.

**Table 3.** Average coefficients of determination (R^2^) from a 5-fold cross-validation for different regression models. The three machine learning models-K-neighbors, linear, and support vector-predict the cannabinoid concentration using data from four different imputation methods: iterative, multiple, k-neighbors, and soft. The best-performing models are bolded. (The R^2^ value for CBDV under K-Neighbors regression using soft imputation values exceeds 1 because of the mean squared error.)

| Cannabinoid | K-Neighbors |  |  |  | Linear |  |  |  | Support Vector |  |  |  |
| --- | --- | --- | --- | --- | --- | --- | --- | --- | --- | --- | --- | --- |
|  | I | M | K | S | I | M | K | S | I | M | K | S |
| CBG | 0.537 | 0.107 | 0.160 | 0.305 | 0.286 | 0.087 | 0.020 | 0.158 | **0.546** | 0.045 | 0.068 | 0.452 |
| THC | 0.634 | 0.304 | 0.353 | 0.597 | 0.403 | 0.403 | 0.323 | 0.357 | 0.526 | 0.443 | 0.408 | **0.688** |
| THCV | 0.309 | 0.155 | 0.068 | **0.628** | 0.091 | 0.029 | 0.041 | 0.062 | 0.011 | 0.012 | 0.008 | 0.065 |
| CBC | 0.395 | 0.169 | 0.047 | 0.236 | **0.459** | 0.321 | 0.053 | 0.150 | 0.367 | 0.241 | 0.077 | 0.361 |
| CBD | 0.811 | 0.515 | 0.515 | 0.747 | 0.528 | 0.454 | 0.385 | 0.403 | **0.815** | 0.562 | 0.542 | 0.627 |
| CBDV | 0.306 | 0.156 | 0.057 | 1.127 | **0.477** | 0.048 | 0.038 | 0.256 | 0.441 | 0.030 | 0.034 | 0.075 |
| CBN | **0.648** | 0.338 | 0.102 | 0.471 | 0.495 | 0.240 | 0.019 | 0.231 | 0.351 | 0.044 | 0.081 | 0.033 |

### Dimensionality reduction

We projected each of the four imputation methods using the UMAP technique (figure 6). Focusing on the four clusters, the UMAP projection clearly segregates high- and low-CBD and THC cultivars. This is an important confirmation that clustering based on the THC and CBD scores alone captured most of the substantive differences found using a non-linear and higher-dimensional technique. Low-THC strains (red and green) and high-THC strains (blue and purple) are likewise spatially proximate in three of the projections. Recall the DBSCAN clustering method used also failed to classify some observed points (blue in Figure 2, brown in Figure 6). Judging by their proximity to the other clusters in these projects, the unclassified points largely belong to the low-THC cultivars (red and green).

**Figure 6.**
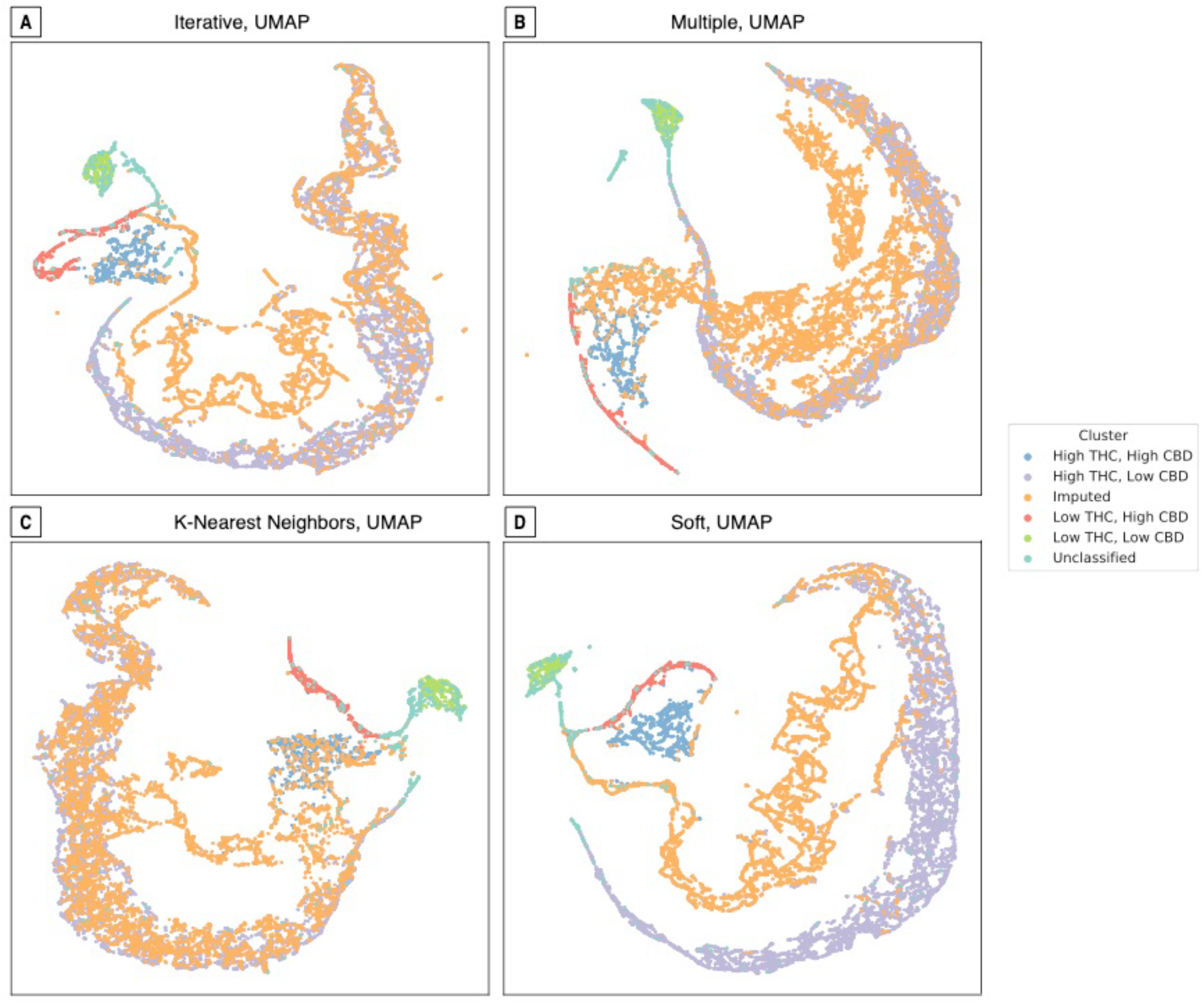
Visualization of imputed data by reducing data with uniform manifold approximation and projection (UMAP).

Cultivars whose CBD and or THC values were imputed are visualized in green (Figure 6). The iterative method (Figure 6A) classified most of these missing values in the high THC, low CBD (orange) cluster, but did identify a “new” grouping closer to the high THC, high CBD (blue) cluster. The projection of the multiple method (Figure 6B) returned at least two distinctive clusters with less overlap with the existing groupings. The projection of the k-nearest neighbors’ method (Figure 6C) unsurprisingly shows a significant overlap with existing values, again mostly with the high-THC and low-CBD and the high-THC and high-CBD clusters. Finally, the values imputed from the soft imputation method (Figure 6D) have minimal overlap with any of the existing clusters and suggests a fifth group somewhere between the high-THC and low-CBD (orange) and the high-THC and high-CBD (blue) clusters.

## Discussion

In this study, we evaluated seven cannabinoids from 17,611 *Cannabis* varieties grown in private US state-markets. Due to the laws and regulations from the different states, we found that many cannabinoids were not tested, and this missing data is not random (Table S1). Therefore, our data consisted mostly of THC and CBD which are the widest-known cannabinoids. We used multiple imputation methods through machine learning techniques to interpolate the missing data. Even though our results suggest cultivars cluster based on their chemotype, and that some imputation methods are better at substituting missing data, due to the non-random absence of particular cannabinoids their behavior could not be accurately imputed. This challenge in the imputation of missing values is particularly difficult for minor cannabinoids such as CBC, CBN, THCV, and CBDV. The results obtained are consistent with the assertion that there is no best method to impute these cannabinoids, and that subsets of the same dataset can behave differently with the same method (ie. Figure 4D, Figure 5C, Table 3).

Our study reaffirms the misnaming of *Cannabis* varieties by the industry (26, 28), since strain identity cannot be predicted according to the clustering groups, even though the clusters are reflective of the chemotype (Figures 3 and S3). Our UMAP projection (Figure 6) segregates high- and low-CBD and THC cultivars, confirming that clustering based on these two cannabinoids captures most of the differences between varieties. Our results also agree with those who have shown that strain name is not indicative of potency or overall chemical composition (35). Due to the misnaming problem we were not able to associate chemotypes to particular groupings. This misnaming problem in the *Cannabis* industry is greatly magnified by the fact that scientist must study the *Cannabis* produced by the federal government studies despite its inferiority in potency and diversity, and the fact that it does not reflect the products from the private markets (19, 30). This is particularly problematic for medical patients who do not understand what they are consuming.

Future work could examine a broader profile of chemotypical data to perform these inferences using ensemble effects which have become a popular analysis method (36). Additionally, as suggested by other studies, terpenoid compounds may be better at clustering the diverse *Cannabis* varieties (36, 37), but perhaps a better approach would be a combined analysis of both cannabinoids and terpenoids (21, 35).

In order to improve the understanding of the *Cannabis* consumed for medical patients, chemotype testing must be made mandatory. However, testing facilities do not have standardized measurement protocols, cannabinoid analysis methods vary widely across laboratories (38), and there are no institutional oversight to validate testing entities or their methodologies. In our study for example, it was hard to discern whether the presence of 0 (zero) for a given cannabinoid was non-detected or not-analyzed, which are two distinct assessments that should be evaluated differently. Given the lack of standardized laboratory methodologies and the lack of supervision of these testing facilities, differences in cannabinoid reporting is expected. However, despite these weaknesses and lack of standardization, some testing facilities do understand the importance and value of accurate chemotyping and see these chemotype tests as more than a service. Thanks to these type of facilities, important cannabinoid and terpenoid research has been achieved (19, 35–37).

The dimensionality reduction visualizations in Figure 6 cannot be compared with each other since they may be randomly transposed and the imputed values “displace” rather than “overlay” the original data. In spite of these limitations, a “good” projection will separate qualitatively distinct data into separate clusters while spatially preserving their latent similarity. Therefore, despite the different tradeoffs from all of our imputation methods, K-nearest (Figure 5C) displayed the best overlap with the original data. However, the coefficients of determination from the regression models (Table 3) did not show this particular method as the most accurate. Linear dimensionality reduction algorithms like PCA (did not perform as well as the non-linear methods like UMAP (Figure 6) and t-SNE. Finally, some cannabinoids are more accurately predicted than others, but no single imputation method or regression model provided consistent performance (Table 3, Figure 4). This could be due to the high amount of missing data (Table 1, Figure S1) but also suggests the need for incorporating theoretically-informed methods like flux balance analysis.

Because THC and CBD have attracted the most research and popular attention of all the cannabinoids and their synthases both compete for the same precursor CBGA, the relationship between these compounds should reveal significant patterns. Our results show that despite the competition for the same precursor, all of these compounds can be present together. Our imputation results preserved relationships such as the negative THC vs. CBD correlation, and positive correlations for CBC vs. CBD, and CBN vs. CBC (Figure 6). In fact, THC has a negative relationship with all other cannabinoids, and CBD and CBC always show a positive relationship (Figure 4). Additionally, CBC is always found in very low levels when compared to THC and CBD (Figure 2 upper half), which could indicate that CBCA synthase is not as good of a competitor for CBGA. The points near each of the axes where the percentage of that given cannabinoid is close to zero (Figure 2, upper half) suggest that those individual cultivars may either bear a synthase gene that is a bad competitor, or a truncated version of the enzyme (12). Given that these three enzymes could be classified as “sloppy enzymes” or “promiscuous” (13–15), it is likely that they could produce each of the other compounds. Therefore, a third hypothesis could be that CBCA synthase is extremely sloppy and produces more of the other cannabinoids, which would explain the larger amounts THC and CBD than CBC.

Some studies suggest that THC has been selected for and that varieties have been bread to increase in THC potency (5, 6). Even though our results do not show this trend, the average THC from our results is much higher than that reported by others (5, 6). THC production is probably a result of gene sequence variation (12), expression levels, and gene copy number variation (7), and there are multiple genes throughout the genome associated with its production (11, 39). However, the expression of these genes could be due to environmental effects, and research suggests that the amount of THC is most likely due to cultivation conditions (35), which have not yet been measured. Additionally, the trends showing an increase on THC through time uses 20 years of data (5, 6), while we are only looking at eight years. This small snapshot of time probably underestimates the whole pattern of increase on THC, which suggests that breeders and growers are selecting for this compound.

## Author Contributions

D.V. and B.K. analyzed the data, wrote the first draft of the manuscript, conceived and lead the project; B.K performed imputation and dimensionality reduction analyzes; R.G., T.B designed, supervised, and provided chemotype data collection. All authors contributed to manuscript preparation.

## Supplementary Information

### Methods

#### Missing data and imputation

**Figure S1.**
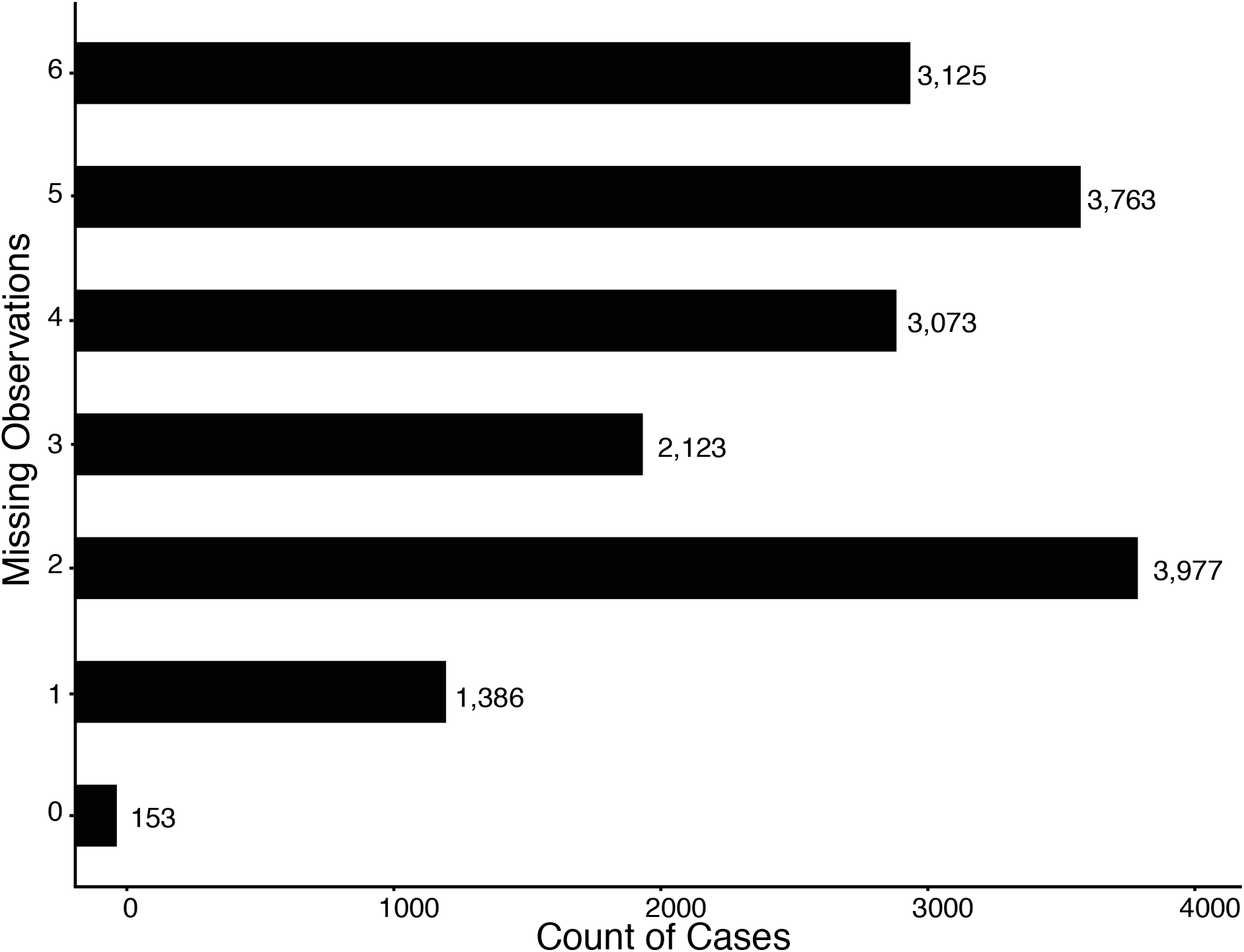
Distribution of cases based on number of missing cannabinoid observations. Only 153 cultivars contain all cannabinoid information, while 1,386 samples miss the information of one of the cannabinoids, 3,977 of two cannabinoids and so on for a total of 17,600 data points analyzed.

The density-based spatial clustering of applications with noise (DBSCAN) algorithm (1) was used to classify points to four clusters using a Euclidian distance metric, a minimum of 75 samples per cluster, and an EPS step size of 0.5.

#### Geographic space analysis

Due to the amount of missing data for CBN, THCV, and CBDV, we did not include these three cannabinoids in the geographic space analysis. We also excluded CBDV and THCB due to the poor understanding of their biochemistry, and CBN because it’s a breakdown product (2–4). Since only 2014 had samples for three states, we analyzed the cannabinoids of these locations only for that year. We had 1,364 samples from California, 936 from Colorado, and 244 from Washington.

We performed a one-way ANOVA for each of four cannabinoids – CBG, CBC, THC, and CBD-as the response variable and state as a factor, with a posterior posthoc analysis to determine cannabinoid level differences by year for each state.

#### Analysis through time

California was the only state that had 16,41 samples from every year for eight years 2011-2018 divided as follows: 361 from 2011; 574 from 2012; 827 from 2013; 1,364 from 2014; 1,113 from 2015; 5 from 2016; 5,727 from 2017; and 6,447 from 2018. Therefore, our analysis of cannabinoid change through time was only performed with samples from this state.

With a one-way ANOVA with each of four cannabinoids – CBG, THC, CBD, and CBC-as the response variable and year as a factor, and with a posterior posthoc analysis we determined whether cannabinoids differ in California in these 8 years.

### Results

#### Geographic space analysis

Despite the disparity in sample size, we find overall differences in cannabinoids for the three states, California, Colorado, and Washington, for the 2014 subset (figure S2). CBG (figure S2A) shows significant overall variation (ANOVA p<0.0001, F=13.09), and the posthoc assessment shows that the three states differ from each other with p-values of less than 0.03. Overall, THC (figure S2B) differs across the three states (ANOVA p<0.01, F = 33.74), and the posthoc analysis reveals that the three states are significantly different from each other. CBD (figure S2C) also differs overall (ANOVA p<0.0001, F = 4.837),however, the pairwise comparisons show significant differences only between California and Colorado. Finally, CBC (figure S2D) varies significantly across the three states (ANOVA p< 0.0001, F=45.08) but the posthoc test shows that California differs from Washington and Colorado, while Washington and Colorado are not significantly different from each other. Essentially, none of the three states have more or less of a particular cannabinoid.

**Figure S2.**
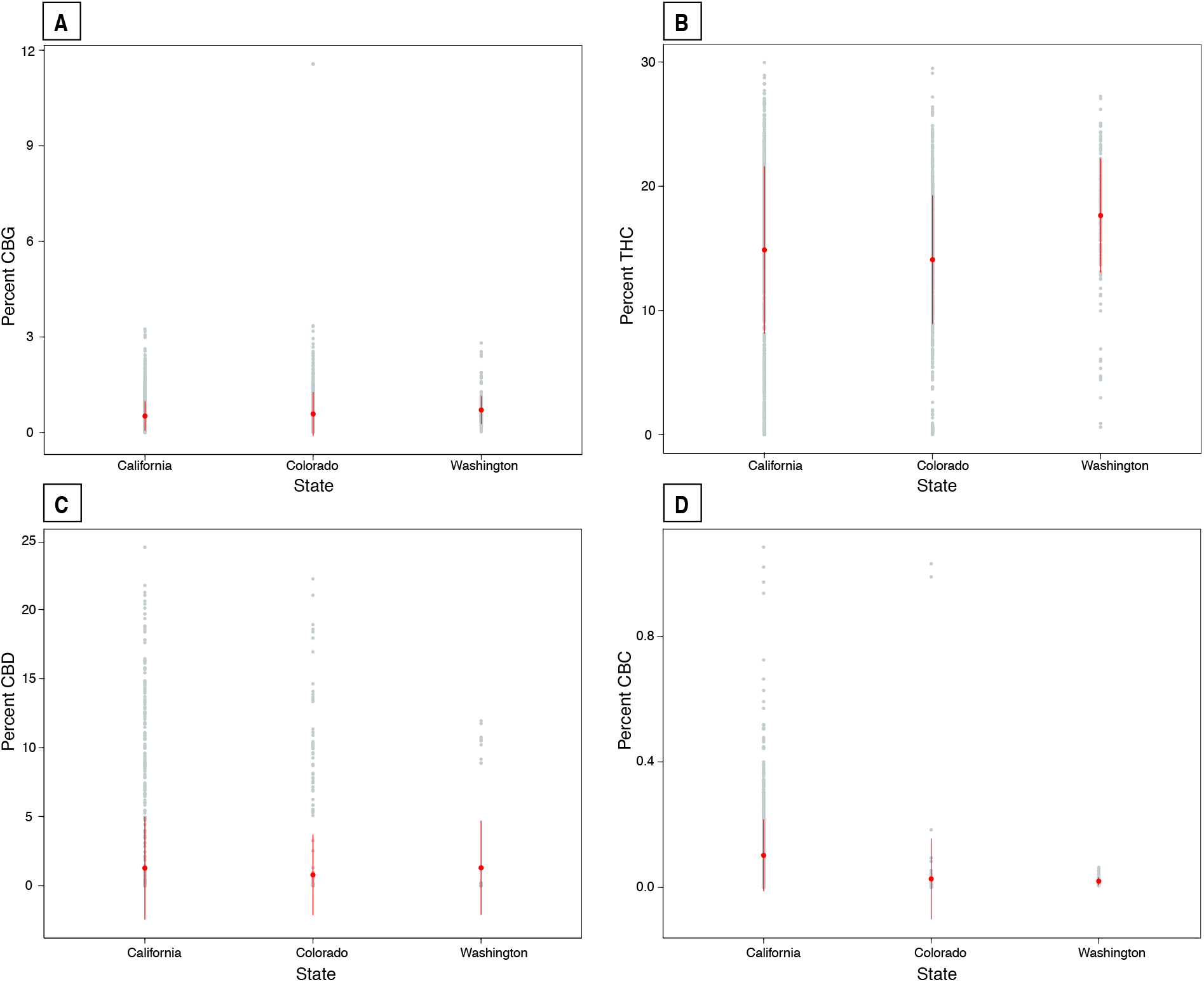
Distribution of four cannabinoids, CBG (A), THC (B), CBD (C), and CBC (D) across three states for 2014. Pairwise comparisons for each panel: **A** California-Colorado p<0.03; California-Washington p<0.0001; Washington-Colorado p<0.01. **B** California-Colorado p<0.01; California-Washington p<0.0001; Washington-Colorado p<0.0001. **C** California-Colorado p<0.01; California-Washington p=NS; Washington-Colorado p=NS. **D** California-Colorado p<0.0001; California-Washington p<0.0001; Washington-Colorado p=NS.

#### Analysis through time

Since California was the only state that had samples from every year for eight years (Table S1), we only included this state for our time analysis for only four of the cannabinoids: CBG, THC, CBD and CBC. With our one way ANOVA with each of the cannabinoids as the response variable and year as a factor, and with a posterior posthoc analysis we found that overall, every single cannabinoid showed significant differences: CBG (F= 234.4, p <0.0001, figure S3A); THC (F = 158.9, p <0.0001, figure S3B); CBD (F= 72.66, p <0.0001, figure S3C); and CBC (F= 5.08, p < 0.03, figure S3D). However, there is not one year where all cannabinoids differ, or there is not one year that differs from all of the other years (Tables S2 and S3). For CBG (Table S2 upper half) or CBD (Table S3 upper half) no year is significantly different from the rest. However, 2015 for THC (Table S2 bottom half) shows significant differences from the rest of the years due to the low levels of the cannabinoid (figure S3B). Similarly, in 2018, CBC (Table S3 bottom half) differs significantly form the other years but due to its high levels (figure S3D). In general, we see no trend for time in increase or decrease of particular cannabinoids in California.

**Figure S3.**
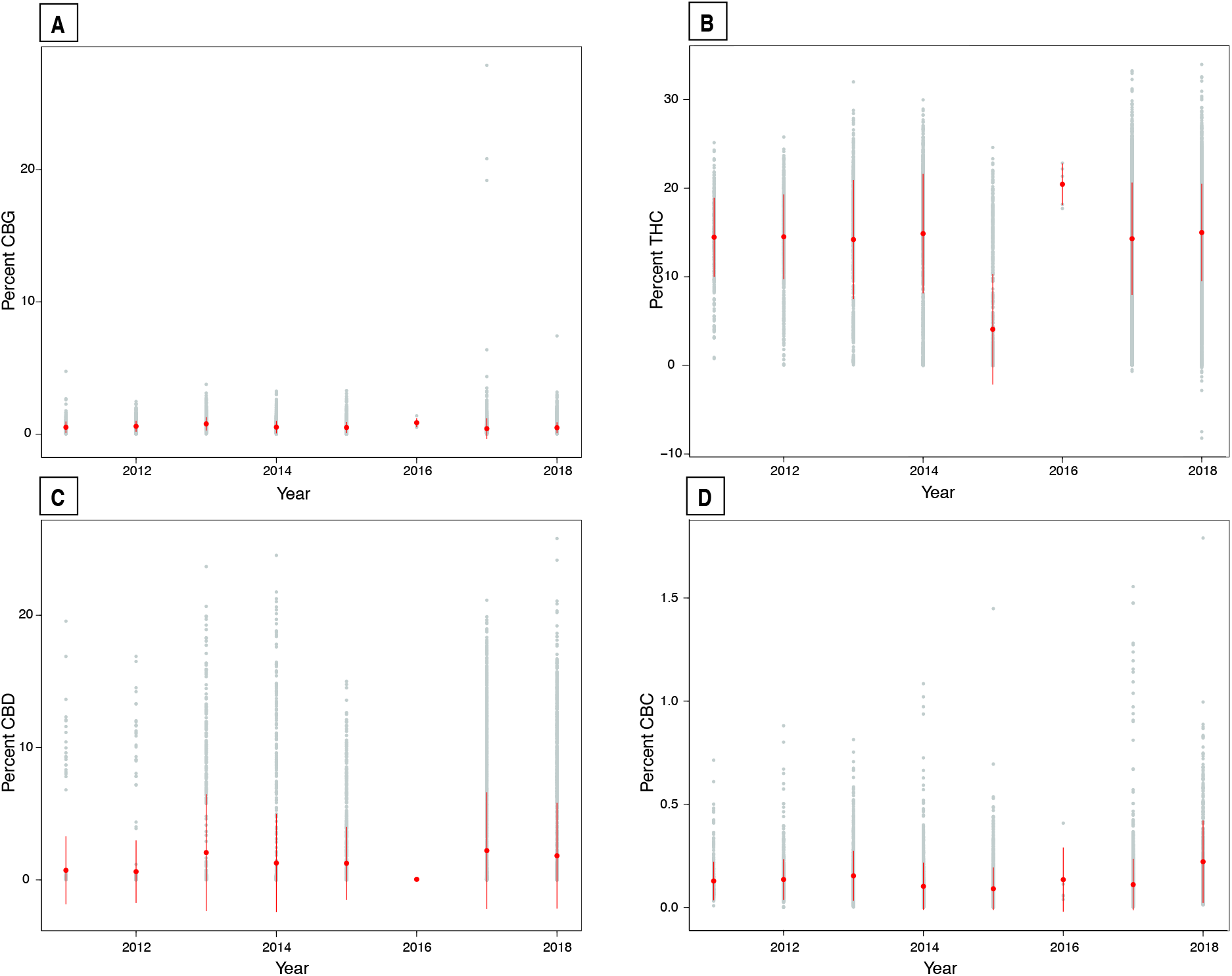
Distribution of four cannabinoids across eight years for the state of California. CBG **(A)** has no year that differs significantly from all other years. 2015 was significantly lower in THC **(B)** production compared to the rest of the years. For CBD **(C)**, no year was significantly different, and for CBC **(D)** 2018 had significantly higher levels when compared to the other seven years. All pair comparisons are given in tables S1 and S2.

**Table S2.** Pairwise comparison between years for CBG (upper triangle) and THC (lower triangle). For both cannabinoids, there is no particular trend in any of the years (Figure S2).

|  |  | CBG California |  |  |  |  |  |  |  |
| --- | --- | --- | --- | --- | --- | --- | --- | --- | --- |
|  |  | 2011 | 2012 | 2013 | 2014 | 2015 | 2016 | 2017 | 2018 |
| THC<br>California | 2011 | - | NS | <0.0001 | NS | NS | NS | <0.0001 | NS |
|  | 2012 | NS | - | <0.0001 | <0.05 | <0.001 | NS | <0.0001 | <0.001 |
|  | 2013 | NS | NS | - | <0.0001 | <0.0001 | NS | <0.0001 | <0.0001 |
|  | 2014 | NS | NS | NS | - | NS | NS | <0.0001 | NS |
|  | 2015 | <0.0001 | <0.0001 | <0.0001 | <0.0001 | - | NS | <0.0001 | NS |
|  | 2016 | NS | NS | NS | NS | <0.0001 | - | NS | NS |
|  | 2017 | NS | NS | NS | NS | <0.0001 | NS | - | <0.0001 |
|  | 2018 | NS | NS | <0.01 | NS | <0.0001 | NS | <0.0001 | - |

**Table S3.** Pairwise comparison between years for CBD (upper triangle) and CBD (lower triangle). For both cannabinoids, there is no particular trend in any of the years (Figure S2).

|  |  | CBD California |  |  |  |  |  |  |  |
| --- | --- | --- | --- | --- | --- | --- | --- | --- | --- |
|  |  | 2011 | 2012 | 2013 | 2014 | 2015 | 2016 | 2017 | 2018 |
| CBC<br>California | 2011 | - | NS | <0.0001 | NS | NS | NS | <0.0001 | <0.0001 |
|  | 2012 | NS | - | <0.0001 | <0.03 | NS | NS | <0.0001 | <0.0001 |
|  | 2013 | <0.05 | NS | - | <0.003 | <0.001 | NS | NS | NS |
|  | 2014 | <0.03 | <0.0001 | <0.0001 | - | NS | NS | <0.0001 | <0.001 |
|  | 2015 | <0.0001 | <0.0001 | <0.0001 | NS | - | NS | <0.0001 | <0.01 |
|  | 2016 | NS | NS | NS | NS | NS | - | NS | NS |
|  | 2017 | NS | <0.001 | <0.0001 | NS | <0.003 | NS | - | <0.01 |
|  | 2018 | <0.0001 | <0.0001 | <0.0001 | <0.0001 | <0.0001 | NS | <0.0001 | - |

#### Dimensionality reduction

Figure S4 is a projection of each of the four imputation methods using the t-SNE technique. The cluster labels using only the CBD and THC relationships in Figure 3 again capture the same clusters in these projections. The relative proximity of the four clusters in these projections appears to be dominated more by similarity in CBD than in THC: the high-CBD, high-THC (blue) is close to the high-CBD, low-THC (red) cluster while the low-CBD, high-THC (orange) and low-CBD, low-THC (purple) clusters are proximate. The iterative method (upper left) projected by t-SNE locates imputed values both in the high-THC, low-CBD (orange) cluster as well as a new cluster closer to the high-THC, high-CBD (blue) cluster. The multiple imputation method (upper right) creates two new distinctive clusters beyond the high-THC, low-CBD (orange) cluster. The k-nearest neighbors (lower left) again locates most of the imputed values inside the high-THC, low-CBD (orange) cluster. Finally, the values imputed from the soft imputation method (lower right) again have minimal overlap with any of the other clusters and form an entirely new cluster between the high-THC and low-CBD (orange) cluster and the high-THC and high-CBD (blue) cluster.

**Figure S4.**
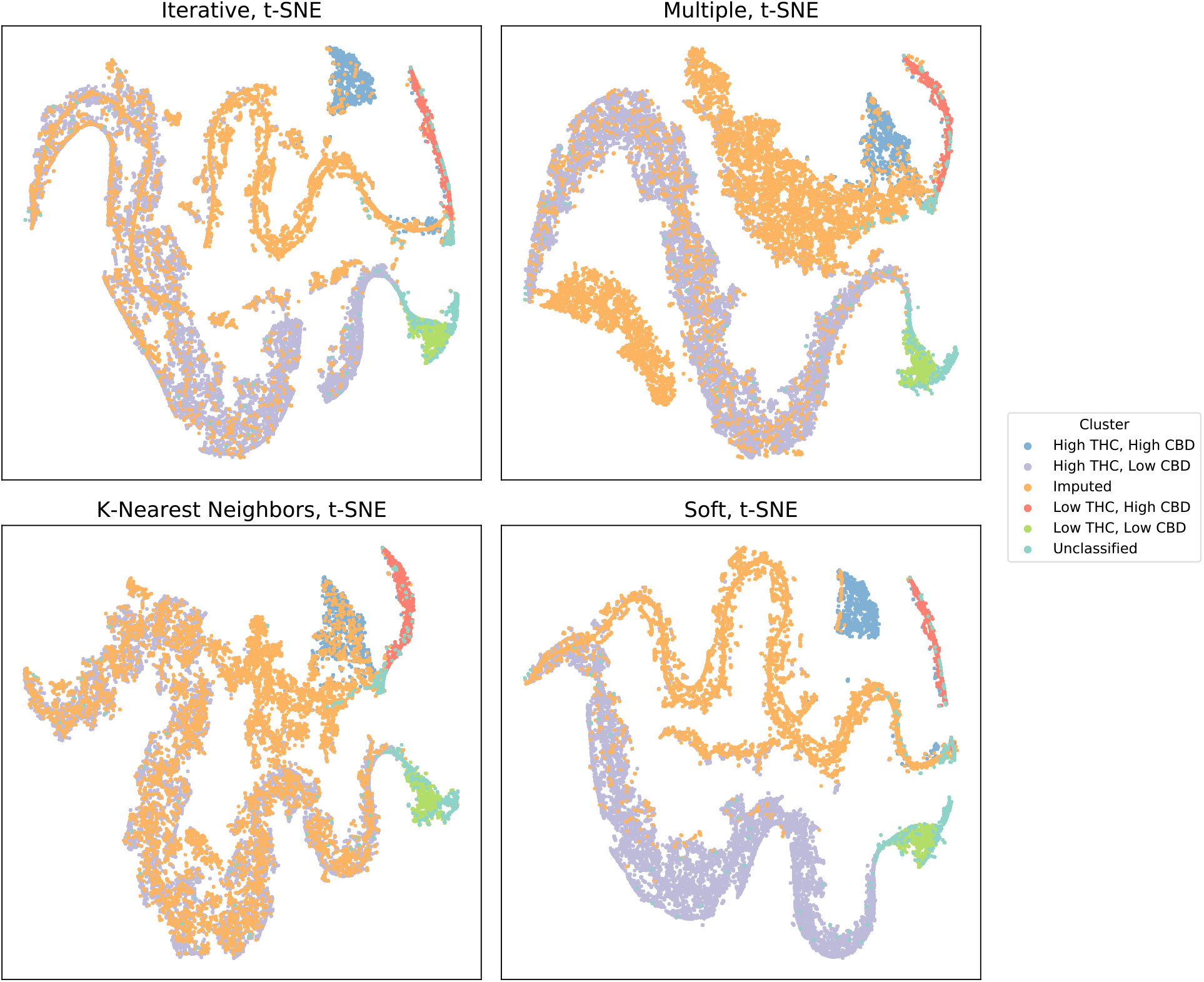
Projection of each of the four imputation methods using the t-SNE. Even though Principal Component Analysis is the most common visualization technique in biology (5), PCA dimensional reduction leads to little or no differentiation between possible groups in these two dimensional mappings (Figure S5).

**Figure S5.**
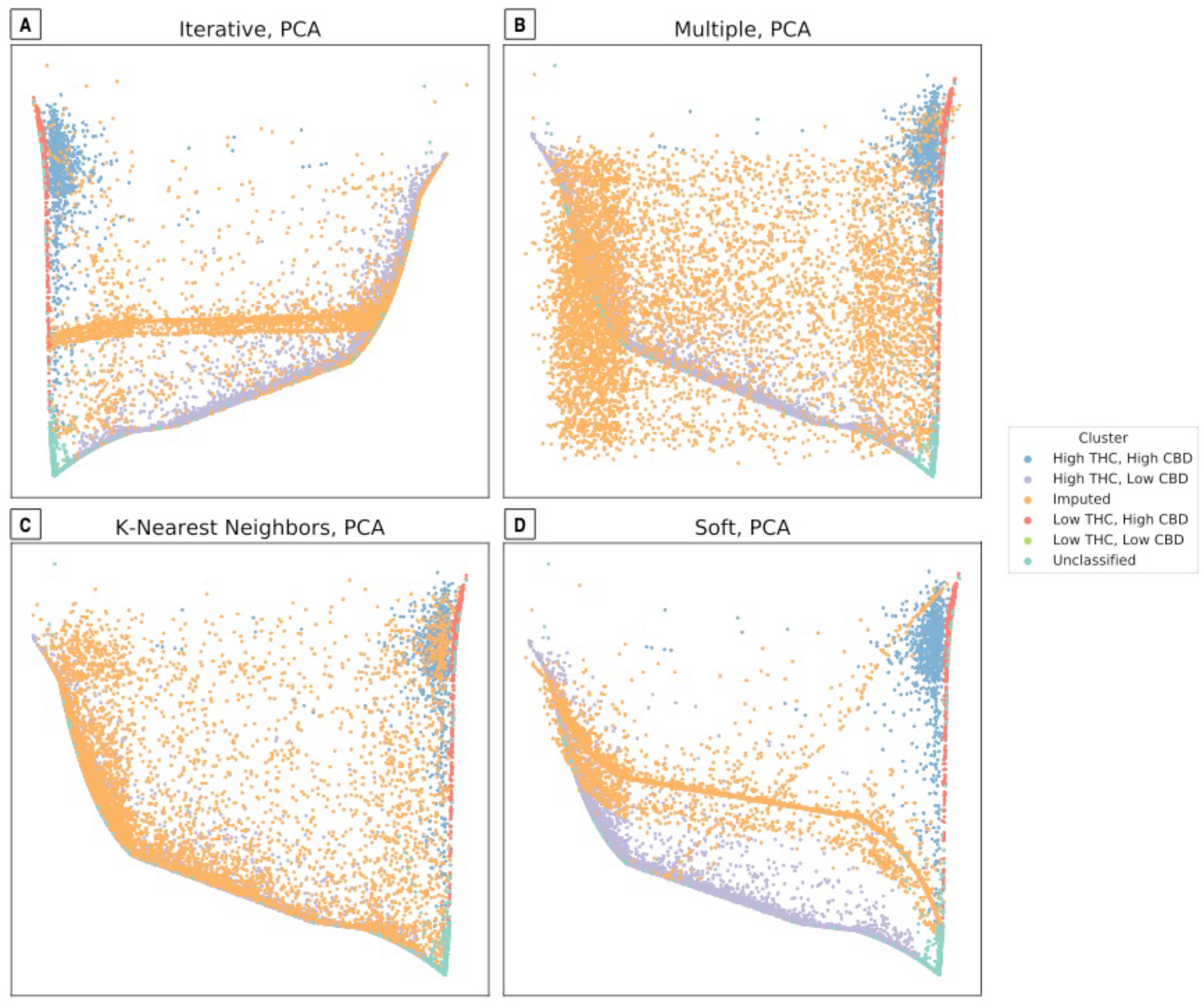
Projection of each of the four imputation methods using PCA

